# Parametric Physics-Based Synthesis of 3D Fluorescence Organoid Images with Exact Ground Truth for Deep Learning Pipeline Development

**DOI:** 10.64898/2026.04.16.719066

**Authors:** Sreenivas Bhattiprolu

**Affiliations:** DigitalSreeni LLC., Antioch CA, USA

**Keywords:** organoid, synthetic image generation, deep learning, fluorescence microscopy, cell segmentation, ground truth, 3D imaging, OME-TIFF, arivis, confocal, cyst, apical-basal polarity

## Abstract

Three-dimensional organoid cultures have emerged as powerful models for studying human tissue biology, disease mechanisms, and drug responses. Fluorescence confocal microscopy of organoids generates complex volumetric image data that is increasingly analyzed using deep learning pipelines for cell segmentation, morphometry, and phenotyping. However, training and benchmarking such pipelines requires large annotated datasets, the manual curation of which is prohibitively expensive and time-consuming. Here we present a parametric, physics-based computational framework for generating synthetic 3D fluorescence organoid images with exact ground-truth cell body and nucleus label masks.

The framework models cell placement using force-directed sphere packing with optional hollow lumen exclusion for cyst-forming organoids, cell morphology using power-diagram (Laguerre) tessellation with apical-basal elongation and surface flattening for polarized epithelial cells, membrane curvature using low-frequency coordinate displacement, nuclear shape using irregular ellipsoid deformation with smooth radial eccentricity direction blending, and optical effects using depth-dependent point-spread function broadening, a physically motivated staining diffusion gradient with residual interior plateau, z-attenuation, haze, shot noise, and channel crosstalk. The necrotic core model uses a three-phenotype nuclear population, pyknotic, ghost, and karyorrhectic, reflecting the histological diversity of real necrotic zones.

Five condition-specific presets are provided, calibrated to published biological measurements and covering PDAC osmotic stress, HMECyst normal and cyst-forming organoids, and a large primary PDAC organoid with a necrotic core. Unlike generative adversarial network approaches, our method requires no training data, produces exact ground truth by construction, and allows systematic and interpretable control over every morphological and optical parameter. The framework is released as open-source Python software with a PyQt5 graphical interface and produces OME-TIFF output compatible with arivis Pro, FIJI, and napari, as well as most other microscopy image analysis software.

## 1. INTRODUCTION

### 1.1 The organoid imaging challenge

Three-dimensional organoid culture systems have transformed biomedical research over the past decade. First described for intestinal epithelium by Sato and colleagues in 2009 [1], organoid technology now encompasses brain, liver, kidney, pancreatic, lung, and cancer-derived systems that recapitulate the structural and functional complexity of native tissue with good fidelity [2]. The morphological diversity of organoids, ranging from compact tumor spheroids to budding intestinal crypts, hollow cyst-forming epithelial structures, and elaborate cerebral organoid architectures, makes them both powerful experimental models and challenging objects for automated image analysis.

Confocal and lightsheet fluorescence microscopy of organoids generates volumetric (3D) image stacks routinely containing hundreds of individual cells per organoid. Quantitative analysis of these stacks such as, counting cells, measuring nuclear morphology, tracking proliferative zones, assessing drug-induced changes, requires robust automated pipelines. Deep learning methods for 3D cell segmentation such as Stardist-3D [3] and Cellpose [4] have matured considerably and provide strong pretrained models that perform well on many standard datasets without requiring additional training. However, as organoids grow larger and more architecturally complex, developing hollow lumens, necrotic cores, or irregular budding morphologies, out-of-the-box models increasingly fail, particularly in regions of dense cell packing, low signal-to-noise ratio, or significant optical attenuation at depth. Adapting these models to new organoid types requires additional annotated training data, and generating that data for 3D fluorescence stacks at single-cell resolution is extraordinarily labor-intensive [6]. More fundamentally, even where pretrained models segment adequately, the field lacks a principled way to benchmark segmentation quality, validate morphometric feature extraction pipelines, or stress-test analysis tools against known ground truth, because real organoid data never comes with exact, pixel-perfect labels.

### 1.2 The case for synthetic organoid images

Synthetic organoid images with exact, mathematically defined ground truth address several distinct needs that real data cannot.

The most direct need is pipeline benchmarking and validation. With real organoid images, there is no way to know whether a segmentation algorithm is correct at the boundary level, whether a morphometric feature extractor handles dense regions accurately, or whether an analysis pipeline breaks silently when organoid morphology changes. Synthetic images, where every cell boundary and nucleus position is known by construction, function as unit tests for image analysis pipelines, enabling objective, quantitative comparison across tools and detection of subtle failure modes such as under-segmentation in packed cores or over-merging at cell-cell interfaces.

A second need is stress-testing against edge cases that are rare or absent in real datasets. Dense nuclei packing, low signal-to-noise conditions, partial imaging depth penetration, strong optical attenuation at depth, and abnormal morphologies such as pyknotic necrotic nuclei can all be generated deliberately and in quantity with a parametric simulator. This is critical for building pipelines that are robust across organoid types and imaging conditions rather than optimized for a single lab’s dataset.

A third need is controlled in silico hypothesis testing. A parametric generator allows systematic variation of one biological parameter at a time, cell size, nuclear deformation, lumen fraction, chromatin texture, staining depth, while holding everything else fixed. This is difficult or expensive to achieve biologically, but straightforward computationally. It enables questions such as: if cells shrink under osmotic stress while maintaining their spatial arrangement, will a given segmentation and feature extraction pipeline correctly detect the volume change? If a cyst forms a hollow lumen, do topology features correctly report the architectural transition?

A fourth need is imaging system simulation. Passing synthetic organoids through physically motivated optical models, depth-dependent point spread function broadening, staining diffusion gradients, shot noise, channel crosstalk, allows pipeline developers to assess how their tools perform across different acquisition conditions and to choose imaging parameters before committing to an experimental protocol.

Finally, annotated training data for deep learning remains a genuine bottleneck for organoid-specific fine-tuning even when pretrained models exist. Synthetic images eliminate labeling effort entirely, providing exact voxel-level 3D ground truth at no annotation cost and enabling pretraining or domain adaptation before fine-tuning on small real datasets. This is especially critical for 3D segmentation models, which rely on spatial continuity across z-slices, something that is difficult to annotate consistently by hand. Critically, because synthetic labels are exact rather than hand-drawn, models trained on them do not inherit the boundary ambiguities that arise when human annotators disagree on cell boundaries in dense or low-contrast regions.

### 1.3 Existing approaches to synthetic biological image generation

Several strategies have been proposed for generating synthetic cell and tissue images to augment or replace real training data. Classical physics-based generators such as the SIMCEP framework for 2D fluorescence cell images [7] and its successors model cell placement, fluorescence intensity, and optics analytically. The CytoPacq framework [8] extended physics-based simulation to 3D, demonstrating that synthetic images can produce training datasets competitive with real annotations.

Generative adversarial networks, including CycleGAN-based unpaired domain transfer [9] and GAN approaches for cell image synthesis [10], can translate between imaging modalities or between synthetic and real image domains. CUT (Contrastive Unpaired Translation) [11] allows a synthetic image to be stylistically transferred to match a target domain without requiring paired data, enabling fine-tuned segmentation models without exact image-label correspondences. Latent diffusion models have more recently shown promise for high-fidelity cell image synthesis [12, 13]. However, all generative deep learning approaches share fundamental limitations when applied to organoid training datasets: they require real training images, they do not natively produce ground-truth annotations, and they offer limited interpretable parameter control over generated morphology.

A third class of approaches uses agent-based simulation to model tissue growth and cell packing directly. CompuCell3D [14] and VirtualLeaf [15] simulate multicellular dynamics using the Cellular Potts Model. These are powerful for studying developmental biology but are computationally expensive for generating large image datasets.

### 1.4 This work

We present a parametric physics-based framework specifically designed for generating synthetic 3D fluorescence organoid images at scale. The key design decisions are: (1) cell placement via force-directed sphere packing with optional hollow lumen exclusion for cyst-forming organoids; (2) cell boundary generation via Laguerre power-diagram tessellation with low-frequency coordinate displacement producing curved biological membranes, unified apical-basal elongation and surface flattening for polarized epithelial cells, and combined radial anisotropy in the distance metric; (3) an explicit physical optics model including depth-dependent PSF broadening, radial staining diffusion with interior residual plateau, z-attenuation, haze, shot noise, and channel crosstalk; (4) a three-phenotype necrotic core model capturing the histological diversity of pyknotic, ghost, and karyorrhectic cells; (5) smooth radial interpolation of nuclear eccentricity direction from random in the core to radially outward at the periphery; and (6) output in OME-TIFF format directly compatible with existing analysis pipelines.

We validate the framework by generating synthetic organoids across three biologically distinct conditions, PDAC spheroids under osmotic stress, normal versus cyst-forming mammary epithelial organoids, and a large heterogeneous primary PDAC organoid, and analyzing their morphometric and spatial topology feature distributions using the same quantitative pipelines applied to real organoid data in published studies. Importantly, the synthetic organoids used for validation are generated from scratch by our framework; we are not attempting to replicate images from any existing dataset. The goal is to show that our generator produces organoids with feature distributions that are plausible and consistent with published biological measurements for the respective organoid types.

## 2 FRAMEWORK DESIGN AND IMPLEMENTATION

### 2.1 Overview

The synthetic organoid generator consists of five sequential computational stages: (1) organoid scaffold and cell placement, (2) cell and nucleus signal rendering, (3) physical optics simulation, (4) OME-TIFF output with embedded metadata, and (5) ground-truth label mask export. The entire pipeline is implemented in Python 3.11 using NumPy, SciPy, and tifffile, with no deep learning dependencies. Generation of a 150 µm diameter organoid (approximately 490 cells, 256×256×160 voxels at 0.414×0.414×1.0 um) requires approximately 90 to 120 seconds on a modern consumer CPU.

### 2.2 Cell placement: force-directed sphere packing

The organoid volume is modeled as a spheroidal boundary with semi-axes derived from the target diameter and a sphericity parameter (1.0 = sphere, 0.75 to 0.95 for typical oblate organoids). Cells are represented as spheres with radii drawn from a lognormal distribution parameterized by zone-specific mean radii (cell_radius_core, cell_radius_periph) and a standard deviation (cell_radius_std). Cells in the inner zone (radial position less than core_fraction) are assigned smaller radii reflecting mechanical confinement; peripheral cells have larger radii consistent with lower confining pressure and greater access to nutrients.

When a hollow lumen is specified (lumen_fraction > 0), an initial exclusion zone is enforced: candidate cell positions whose normalized radial distance from the organoid center is less than lumen_fraction are rejected during seeding. During the subsequent force-directed relaxation, any cell that drifts inside the lumen boundary receives a centrifugal repulsion force scaled by boundary_strength x 1.5, preventing cells from settling back into the void. The lumen renders as background fluorescence by construction.

The force-directed relaxation applies repulsion forces between overlapping cells and boundary forces to cells outside the organoid ellipsoid, integrated with damped Euler steps.

*Repulsion force between overlapping cells i and j:*

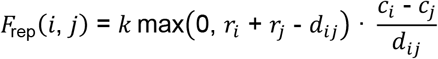

where k = repulsion_strength, *r*_*i*_ is the radius of cell *i, d*_*i*_ is the inter-center distance, and *c*_*i*_ is the center of cell *i*.

*Lumen repulsion force (active when lumen_fraction > 0):*

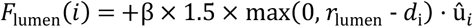

where β = boundary_strength, *r*_lumen_ is the lumen exclusion radius, d_i_ is the distance of cell *i* from the organoid center, and û_i_ is the unit outward radial direction.

This algorithm converges in 60 to 150 iterations for typical organoid sizes.

### 2.3 Cell body rendering: Laguerre power-diagram tessellation with apical-basal polarity

Each voxel is assigned to the cell that minimizes a unified anisotropic power distance. The distance metric is parameterized by a per-cell radial scale factor (cell_apical_scale) combining three effects along the outward radial axis.

*Radial scale factor combining compression, apical elongation, and surface flattening:*

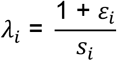

where ε_i_ = comp_fac_i_ is the radial compression factor and *s*_*i*_= cell_apical_scalei (*s*_*i*_> 1: columnar elongation; *s*_*i*_< 1: squamous flattening).

*Anisotropic power distance for Laguerre tessellation:*

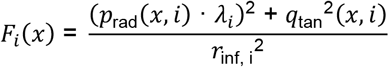

where *p*_rad_(*x, i*) is the radial displacement component from cell center *c*_*i*_, *q*_tan_(*x, i*) is the tangential component, and *r*_inf_,_i_= *r*_*i*_(1 + ρ_i_) is the inflated cell radius with pressure factor ρ_i_.

The comp_fac term from radial_compression flattens core cells along the radial axis. The cell_apical_scale encodes two additional effects:

- **apical_elongation** (> 0): stretches peripheral cell territories radially. A t-squared ramp increases apical_scale from 1.0 at core_fraction to 1 + apical_elongation at the organoid surface. When cell_apical_scale > 1, cells claim more territory radially, producing columnar shapes taller radially than tangentially.
- **surface_flattening** (> 0): compresses the outermost cells radially. Applied only in the outermost 15% of the organoid radius with a linear ramp, reducing flat_scale toward 1 - surface_flattening at the surface. When cell_apical_scale < 1, cells claim less radial territory, producing squamous-like flattening against the ECM/air interface.

Both effects act multiplicatively: cell_apical_scale = apical_scale x flat_scale. When both parameters are 0.0, cell_apical_scale = 1.0 exactly and the formula reduces to the previous behavior, preserving full backward compatibility with all existing presets.

### 2.4 Membrane curvature: coordinate displacement field

The Voronoi cell boundaries produced by a pure power-diagram are mathematically flat faces. We introduce a low-frequency 3D displacement field that perturbs query coordinates before the power-diagram competition, causing effective boundaries to follow smoothly curved paths:

*Low-frequency coordinate displacement before power-diagram competition:*

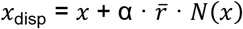

where α = membrane_bend_amplitude, 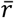 is the mean cell radius, and *N*(*x*) is a unit-normalized low-frequency 3D noise field at spatial scale 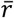. The physical clip check uses original coordinates *x* (not *x*_disp_) to prevent cells extending beyond their physical radius.

### 2.5 Nucleus rendering with smooth eccentricity direction blending

For each cell, a nucleus center is computed with an eccentricity offset whose direction transitions smoothly from random in core cells to radially outward in peripheral cells. The updated model uses a t-squared blend weight.

*t-squared blend weight for smooth eccentricity direction transition:*

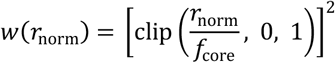

where *r*_norm_ is the normalized radial position (0 = organoid center, 1 = surface) and *f*_core_ = core_fraction. At w = 0 the nucleus eccentricity direction is purely random; at w = 1 it is radially outward.

*Smooth blend of nucleus eccentricity direction:*

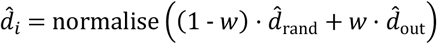

Where 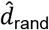 is a unit random direction and 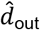 is the unit outward radial direction.

At *w*= 0 (organoid center) the direction is purely random; at the surface it is strongly outward. This produces a realistic gradient of nuclear positioning matching the gradual onset of epithelial polarity observed in organoid cross-sections. The magnitude of eccentricity is independently interpolated between nucleus_ecc_core and nucleus_ecc_periph.

Nucleus shape is modeled as a triaxial ellipsoid with a fractal surface deformation

*Nucleus shape: triaxial ellipsoid with fractal surface deformation:*

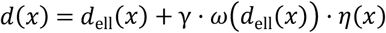

where *d*_ell_(*x*) is the triaxial ellipsoid distance, γ = nucleus irregularity, ω(d) = clip(d/1.1, 0, 1)^2^ concentrates deformation near the nucleus surface, and η(x) is a three-octave fractal noise field seeded per cell.

### 2.6 Actin/membrane signal rendering

The actin channel represents phalloidin staining concentrated at the cortical actin cytoskeleton at cell-cell interfaces and the outer cell cortex. Any voxel whose face-adjacent neighbor belongs to a different cell is marked as membrane. Outer cell surfaces facing the organoid exterior receive identical treatment to inner interfaces. A faint cytoplasm fill (actin_cytoplasm_frac x membrane_brightness) is added within each cell body using a spatially heterogeneous low-frequency noise field. Tricellular junctions, voxels at the intersection of three or more cell boundaries, receive a brightness boost of 1.35x, reflecting known enrichment of junction complex proteins at these sites.

### 2.7 Necrotic core simulation with three-phenotype nuclear population

Organoids larger than approximately 150 µm in diameter develop a hypoxic/necrotic core due to diffusion limits on oxygen and nutrients. When the necrotic_core parameter is enabled, cells in the innermost necrotic_fraction of the organoid radius are assigned to the necrotic zone. Thirty percent of these cells are randomly excluded from rendering entirely, producing the dark voids characteristic of necrotic cores. Actin signal is set to zero for all necrotic-zone cells, representing membrane breakdown.

The remaining necrotic-zone cells are assigned to one of three nuclear phenotypes, deterministically seeded by cell_id for reproducibility:

- **Pyknotic (65%)**: condensed chromatin. Nucleus radius scaled to uniform(0.50, 0.72) x normal, DAPI brightness multiplied by uniform(0.85, 1.15) x necrotic_dapi_boost. Bright, small nuclei characteristic of apoptotic condensation.
- **Ghost cells (20%)**: nucleus largely dissolved. Radius scaled to uniform(0.88, 1.00) x normal, brightness multiplied by uniform(0.15, 0.35). Dim, nearly full-size nuclei representing late-stage necrotic dissolution (karyolysis).
- **Karyorrhectic (15%)**: intermediate fragmentation. Radius scaled to uniform(0.60, 0.80) x, brightness multiplied by uniform(0.50, 0.85) x necrotic_dapi_boost. Partially condensed nuclei at an intermediate stage.

The necrotic_dapi_boost parameter controls the brightness of pyknotic and karyorrhectic cells; ghost cell dimness is fixed independently of this parameter.

### 2.8 Staining diffusion gradient with interior residual plateau

Fluorescent dyes and phalloidin penetrate organoids by diffusion from the outer surface. After rendering, both DAPI and actin channels are multiplied by a radial staining attenuation map computed using the 3D Euclidean distance transform from the organoid surface:

*Radial staining attenuation with interior residual plateau:*

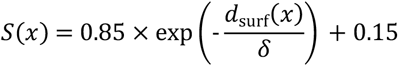

where *d*_surf_(*x*) is the 3D Euclidean distance from voxel *x* to the nearest organoid surface voxel and δ = staining_depth_um. The 0.15 plateau reflects incomplete dye washout in uncleared organoids.

The exponential component represents the primary penetration gradient. The 0.15 additive residual plateau reflects incomplete washout of phalloidin and DAPI in uncleared organoid samples, real confocal images of thick uncleared spheroids consistently show a dim but non-zero interior signal rather than a complete falloff to background. Setting staining_depth_um (δ) = 9999 combined with cleared = True gives effectively uniform staining appropriate for tissue-cleared samples.

### 2.9 Depth-dependent PSF broadening

Light scattering through biological tissue progressively broadens the effective point spread function with increasing imaging depth. We model this by processing the volume in Z-slabs with a depth-varying Gaussian blur.

*Depth-varying lateral PSF sigma:*

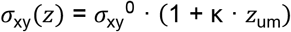

where 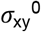 is the diffraction-limited lateral PSF sigma at the coverslip, κ = scatter_increase_rate (per μm), and z_um_ is the imaging depth in μm.

The slab height is set to Z/16, with a margin of 3x sigma_z voxels to prevent edge artifacts at slab boundaries. This produces images where deep slices are genuinely fuzzier, not merely dimmer. Typical values: 0.0 for tissue-cleared or lightsheet samples, 0.003 for confocal imaging of uncleared 100 µm organoids, 0.008 for widefield imaging of thick uncleared samples.

### 2.10 Additional optics: attenuation, haze, noise, and crosstalk

Z-attenuation models overall intensity falloff with depth due to absorption and incomplete confocal rejection:

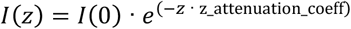

This is applied after PSF blurring, matching the physical acquisition sequence. Out-of-focus haze is modeled as a heavily blurred copy of the volume added at amplitude haze_amplitude. Shot noise is modeled as Poisson-distributed photon noise; read noise is added as zero-mean Gaussian with standard deviation read_noise_std.

Channel crosstalk applies a bidirectional fraction:

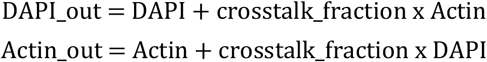

### 2.11 Full parameter reference

A complete parameter reference including type, default value, valid range, and biological interpretation for all parameters is provided in Table 1 at the end of this manuscript.

**Table 1.** Complete parameter reference for the synthetic organoid generator.

| Parameter | Section | Type | Default | Range | Description |
| --- | --- | --- | --- | --- | --- |
| diameter_um | shape | float | 150 | 50-300 | Overall organoid diameter. Sets approximate cell count at given cell size. |
| diameter_std_um | shape | float | 10 | 0-30 | Run-to-run size variation. Set to 0 for reproducible exact size. |
| sphericity | shape | float | 0.92 | 0.75-1.0 | 1.0 = perfect sphere. Lower = flattened along Z axis (oblate). Values below 0.85 give a clearly disc-like shape. |
| cell_radius_core | cells | float (um) | 6.5 | 4-12 | Radius of inner core cells. Below 4 $\mu\text{m}$ nuclei become unresolvable at typical voxel sizes. |
| cell_radius_periph | cells | float (um) | 10.0 | 6-16 | Radius of peripheral cells. Ratio to core controls steepness of the radial gradient. |
| cell_radius_std | cells | float (um) | 0.6 | 0-2 | Cell-to-cell size variation (lognormal sigma). Higher = more heterogeneous population. |
| core_fraction | cells | float | 0.55 | 0.3-0.75 | Fraction of organoid radius defined as the core zone. Also sets the ramp start for apical_elongation. |
| nc_ratio_core | cells | float | 0.72 | 0.50-0.82 | Nucleus-to-cell radius ratio in core. Higher = nuclei fill more of each cell. Volume ratio = this value cubed. |
| nc_ratio_periph | cells | float | 0.70 | 0.48-0.80 | NC ratio at periphery. Usually slightly lower than core. |
| elongation_core | cells | float | 0.88 | 0.75-1.0 | Nucleus short/long axis ratio in core. 1.0 = sphere. Floor is 0.75. |
| elongation_periph | cells | float | 0.78 | 0.65-1.0 | Nucleus elongation at periphery. Lower = more elongated (columnar epithelium). |
| nucleus_ecc_core | cells | float | 0.08 | 0.0-0.20 | Nuclear offset in core cells as fraction of r_cell. Direction is random (no polarity). |
| nucleus_ecc_periph | cells | float | 0.22 | 0.0-0.35 | Nuclear offset in peripheral cells. Direction ramps from random to radially outward via t-squared blend. |
| nucleus_irregularity | cells | float | 0.25 | 0.0-0.40 | Surface deformation of nucleus. 0 = perfect ellipsoid, 0.35 = strong kidney/bean shape (high-grade carcinoma). |
| pressure_core | cells | float | 0.18 | 0.0-0.40 | Compression of core cells. Higher = more compressed, flatter contact faces. Tumor spheroids: 0.22-0.35. |
| pressure_periph | cells | float | 0.06 | 0.0-0.20 | Compression of peripheral cells. Usually 3 to 4x lower than core. |
| radial_compression | cells | float | 0.15 | 0.0-0.30 | Oblate flattening of core cells along the radial axis. 0 = sphere, 0.25 = noticeably oblate. |
| lumen_fraction | cells | float | 0.0 | 0.0-0.80 | Hollow lumen radius as fraction of organoid radius. 0 = solid spheroid. 0.78 = thin polarised shell (cyst mode). |
| apical_elongation | cells | float | 0.0 | 0.0-0.60 | Radial stretch of peripheral cell territories. 0 = isotropic. 0.25 = mild columnar. 0.50 = clearly columnar crypt. Ramps from core_fraction to surface via t-squared weight. |
| surface_flattening | cells | float | 0.0 | 0.0-0.60 | Radial compression of the outermost cells (outermost 15% of radius). 0 = none. 0.30 = visible squamous outer layer. Mimics cells pressed against ECM or air interface. |
| necrotic_core | cells | bool | false | true/false | Enable the necrotic zone. Cells in the innermost necrotic_fraction receive three-phenotype DAPI treatment and zero actin. |
| necrotic_fraction | cells | float | 0.25 | 0.05-0.50 | Inner radius fraction assigned to the necrotic zone when necrotic_core is enabled. |
| necrotic_dapi_boost | cells | float | 1.6 | 1.0-3.0 | Brightness multiplier for pyknotic and karyorrhectic nuclei. Ghost cell dimness is fixed independently. |
| n_iterations | packing | int | 120 | 60-300 | Force-directed relaxation steps. More = tighter packing, slower generation. |
| repulsion_strength | packing | float | 0.45 | 0.2-0.8 | Repulsion force between overlapping cells. Higher = more uniform spacing. |
| damping | packing | float | 0.55 | 0.3-0.8 | Velocity damping per relaxation step. Lower = faster settling, higher = more stable. |
| boundary_strength | packing | float | 1.2 | 0.5-2.0 | Force keeping cells inside the organoid boundary. Also scales lumen repulsion (x1.5) when lumen_fraction > 0. |
| target_overlap | packing | float | 0.02 | 0.0-0.10 | Target fractional overlap between adjacent cells after relaxation. Controls tightness of final packing. |
| nucleus_texture_scale | texture | float | 0.20 | 0.05-0.50 | Spatial frequency scale of the chromatin texture noise field. Lower = coarser texture, higher = finer granularity. |
| nucleus_texture_octaves | texture | int | 3 | 1-5 | Number of fractal octaves in the chromatin noise field. More octaves = more complex multi-scale texture. |
| nucleus_texture_amplitude | texture | float | 0.35 | 0.0-0.60 | Strength of chromatin texture inside nuclei. 0 = uniform fill, 0.6 = strongly granular. |
| membrane_thickness_vox | texture | float | 2.0 | 1.0-4.0 | Actin membrane thickness in voxels. Higher = thicker, more prominent membranes. |
| actin_cytoplasm_frac | texture | float | 0.08 | 0.0-0.25 | Fraction of membrane brightness in the cytoplasm. Higher = more filled-in cells. |
| membrane_bend_amplitude | texture | float | 0.20 | 0.0-0.35 | Low-frequency noise displacement applied before the power-diagram competition. 0 = flat Voronoi faces. 0.20 = biologically natural curves. Critical for avoiding straight-edge artifacts. |
| dapi_intensity_sigma | texture | float | 0.18 | 0.0-0.40 | Cell-to-cell DAPI brightness variation (lognormal sigma). Higher = more heterogeneous staining. |
| actin_intensity_sigma | texture | float | 0.28 | 0.0-0.45 | Cell-to-cell actin brightness variation. |
| dapi_brightness_core | texture | float | 0.70 | 0.3-1.0 | Relative DAPI brightness scaling for core cells. Values below 1.0 produce a dimmer interior consistent with staining gradient effects. |
| dapi_brightness_periph | texture | float | 1.00 | 0.5-1.0 | Relative DAPI brightness scaling for peripheral cells. Typically 1.0 (full brightness at organoid surface). |
| actin_brightness_core | texture | float | 0.55 | 0.2-1.0 | Relative actin brightness scaling for core cells. Reflects reduced membrane signal in compressed inner cells. |
| actin_brightness_periph | texture | float | 1.00 | 0.5-1.0 | Relative actin brightness scaling for peripheral cells. Typically 1.0. |
| heterochromatin_fraction | texture | float | 0.25 | 0.0-1.0 | Fraction of chromatin texture represented as discrete heterochromatin blobs versus smooth euchromatin background. High values = granular punctate texture (open/active chromatin). Low values = uniform compacted texture. Used to differentiate isotonic (0.30) vs hypertonic (0.05) chromatin states. |
| psf_sigma_xy_um | optics | float (um) | 0.25 | 0.15-0.50 | Lateral PSF Gaussian sigma. Match to objective NA: 0.20 = high NA confocal, 0.40 = widefield. |
| psf_sigma_z_um | optics | float (um) | 1.20 | 0.5-3.0 | Axial PSF sigma. Always larger than XY: 0.65 = lightsheet, 1.2 = confocal, 2.0 = widefield. |
| z_attenuation_coeff | optics | float | 0.005 | 0.001-0.015 | Overall intensity falloff with depth. Match to sample thickness and clearing status. |
| staining_depth_um | optics | float | 35.0 | 5-9999 | Half-penetration depth of dye from organoid surface. Used in $0.85 \times \exp(-d/\text{depth}) + 0.15$ model. 9999 = uniform staining. |
| scatter_increase_rate | optics | float | 0.005 | 0.0-0.012 | PSF broadening per $\mu\text{m}$ depth. 0 = cleared, 0.003 = confocal uncleared, 0.008 = widefield thick sample. |
| haze_sigma_um | optics | float (um) | 10.0 | 5.0-30.0 | Spatial scale of the out-of-focus haze blur kernel in um. Larger = broader diffuse glow from adjacent planes. |
| haze_amplitude | optics | float | 0.10 | 0.0-0.25 | Out-of-focus glow from adjacent planes. Higher = more diffuse background. |
| shot_noise_scale | optics | float | 0.05 | 0.01-0.12 | Poisson photon noise scale. Higher = grainier, lower SNR. |
| read_noise_std | optics | float | 0.012 | 0.003-0.030 | Gaussian detector readout noise. Adds a constant noise floor independent of signal level. |
| background_level | optics | float | 0.04 | 0.01-0.10 | Constant background offset. Higher = brighter background as in uncleared samples. |
| crosstalk_fraction | optics | float | 0.04 | 0.0-0.15 | Bidirectional channel bleed-through between DAPI and actin. |
| cleared | optics | bool | false | true/false | Tissue-clearing flag. Disables staining gradient and depth-dependent PSF broadening. Use the apply_clearing() convenience method. |

### 2.12 Output format

All synthetic images are saved as OME-TIFF files with embedded OME-XML metadata specifying voxel physical dimensions and channel names. Three files are produced per run: the two-channel fluorescence image (DAPI + Actin), a cell body integer label mask (uint16, one unique integer per cell), and a nucleus label mask (same cell IDs, nucleus voxels only). Nucleus and cell label IDs are identical, allowing direct linkage without a lookup table. Files are auto-named with preset name, random seed, and timestamp to prevent overwriting. Output is directly loadable in arivis Pro, FIJI/ImageJ (Bio-Formats), napari, and 3DCellScope [5].

## 3 BIOLOGICAL PARAMETERISATION OF ORGANOID TYPE PRESETS

### 3.1 Parameter grounding philosophy

We partition parameters into three categories by evidential basis. Category A parameters are directly grounded in published quantitative measurements. Category B parameters are qualitatively motivated by published biology but lack precise quantitative data and represent reasoned estimates. Category C parameters are purely algorithmic with no biological interpretation and should be set for numerical stability.

### 3.2 Category A - Literature-grounded parameters

The nucleus-to-cytoplasm (NC) ratio is among the most rigorously measured parameters in cell biology, serving as a quantitative diagnostic criterion in cytopathology. Imaging flow cytometry measurements across four cancer cell lines consistently find NC ratios (linear diameter) of 0.60 to 0.70, with the non-malignant breast epithelial line MCF-10A measuring 0.53 [16]. Our presets use cancer values of 0.70 to 0.78 and normal epithelium values of 0.60 to 0.68.

Cell size ranges are well established. Hepatocytes: 20 to 30 µm diameter [17]; typical carcinoma cells: 15 to 25 µm; intestinal columnar epithelial cells: 8 to 15 µm diameter, 20 to 40 µm height. Nuclear pleomorphism is a formal grading criterion in tumor pathology (WHO Classification of Tumors) [18]: smooth round nuclei (Grade 1), mild irregularity (Grade 2), marked pleomorphism and multilobular nuclei (Grade 3). Our nucleus_irregularity parameter (0.10 to 0.40) maps directly to this histological grade scale.

The necrotic core is a well-documented feature of spheroids above approximately 150 µm diameter [19]. The inner core exhibits pyknotic nuclei and absent actin signal [20]. The three-phenotype nuclear population model reflects established necrosis pathology: pyknosis and karyorrhexis are recognized histological stages preceding karyolysis, corresponding to our ghost cell phenotype.

### 3.3 Category A - Literature-grounded parameters

Pressure parameters model the mechanical environment of cells in packed tissue. Published measurements show that tumor spheroids develop internal hydrostatic compressive stress of approximately 1 to 10 kPa [21, 22]. Brain tissue has Young’s modulus of 0.1 to 1 kPa [23]. These relative orderings motivate our pressure parameter choices, but the absolute values should be calibrated against morphometric measurements from real images.

The staining depth parameter is motivated by diffusion physics. For small molecule dyes such as DAPI and phalloidin conjugates, penetration length L is approximately 30 to 80 μm for typical overnight staining protocols. For antibodies (MW ~150 kDa), L is approximately 10 to 30 μm. The residual 0.15 plateau in the staining model is an empirical estimate based on observed dim interior signal in published confocal cross-sections of uncleared organoids.

The apical_elongation and surface_flattening parameters have no published quantitative equivalent. They are specific to our power-diagram-based cell territory model. Values in the calibrated presets were tuned visually by comparing generated center-plane slices against confocal images of the respective organoid types.

### 3.4 Category C - Algorithmic parameters

Packing parameters (n_iterations, repulsion_strength, damping, boundary_strength) control numerical convergence of the sphere packing algorithm and have no biological interpretation. Similarly, the noise frequencies and octave counts used in nucleus texture and membrane curvature generation are aesthetic choices producing visually realistic variation; they do not correspond to specific biological length scales.

### 3.5 Condition-specific calibrated presets

Five presets target specific organoid conditions were drawn from the published dataset of Ong et al. (2025) [5]. Preset parameters were calibrated by applying our feature extraction pipeline to the open-access per-cell data published by Ong et al., extracting spatial topology and morphology distributions for each condition, and tuning preset parameters until the synthetic feature distributions matched the real ones. Key calibration targets included crystal distance (22.7 μm for HMECyst control, 17.3 μm for cyst), nuclear density (94,869 vs 120,329 nuclei/mm3), Ripley ratio (1.685 vs 2.721), and cell volume (~2,300 μm^3^ for PDAC). The imaging parameters (psf_sigma_xy = 0.22 μm, psf_sigma_z = 0.95 μm, voxel_xy = 0.345 μm, z_step = 1.0 um) correspond to a Nikon Ti2-E AX confocal with a 25x Plan Apo Lambda S silicone 1.05 NA objective, NucBlue + Alexa Fluor 647 Phalloidin staining, Fluoromount G mounting, matching the imaging conditions described in Ong et al. (2025) [5]. This ensures that the same feature definitions and analysis pipeline are used for both parameter calibration and synthetic validation, making comparisons between synthetic and real organoids directly interpretable.

Calibrated parameter values for all five presets are summarized in Table 2 at the end of this manuscript.

**Table 2.** Condition-specific calibrated presets.

| Preset | Diam. | ~Cells | Core to Periph (um) | Pressure c/p | NC ratio | Irreg. | Character |
| --- | --- | --- | --- | --- | --- | --- | --- |
| pdac_isotonic | 170 | ~640 | 7.0 to 9.5 | 0.18/0.06 | 0.72/0.68 | 0.26 | PDAC spheroid, isotonic control. nucleus_texture_amplitude=0.38. Calibrated to cell volume ~2300 um<sup>3</sup>, elongation 1.40 (major/minor). |
| pdac_hypertonic | 165 | ~610 | 7.0 to 9.5 | 0.22/0.08 | 0.72/0.68 | 0.30 | PDAC spheroid, osmotic stress. nucleus_texture_amplitude=0.38. Slight nuclear volume decrease vs isotonic. Chromatin more uniformly distributed. Cell shape and topology preserved. |
| hmecyst_control | 195 | ~260 | 12.0 to 14.5 | 0.12/0.05 | 0.65/0.62 | 0.14 | HMECyst normal epithelial. Calibrated to nuclear density 94,869/mm<sup>3</sup> and crystal distance 22.7 um. Solid spheroid, disordered packing (Ripley ratio ~1.7). |
| hmecyst_cyst | 190 | ~40-60 | 10.5 to 13.0 | 0.08/0.20 | 0.65/0.63 | 0.12 | HMECyst cyst-forming. lumen_fraction=0.78. Calibrated to Ripley ratio 2.721 (crystal-like shell). apical_elongation=0.15, surface_flattening=0.15. |
| pdac_large_clustering | 200 | ~1000 | 6.5 to 10.0 | 0.28/0.09 | 0.74/0.70 | 0.34 | Large primary PDAC. necrotic_core=true, necrotic_fraction=0.28. Continuous radial gradient; topology and size features dominate cluster separation. |

**Table 3.** Physics-Based vs Deep Learning synthesis.

| Parametric Physics-Based (This Work) | GAN / Deep Learning Synthesis |
| --- | --- |
| No training data required - generates from parameters alone | Requires large corpus of real annotated images (circular dependency) |
| Exact ground truth by construction - cell and nucleus labels are mathematically precise | Ground truth must be estimated post-hoc or via paired registration, introducing error |
| Full interpretable parameter control - NC ratio, pressure, clearing, staining depth, apical polarity all explicit | No direct parameter control - generated distribution constrained by training data |
| Systematic parameter sweeps - generate 100 images at staining_depth = 10, 20 ... 100 $\mu\text{m}$ | Cannot generate controlled parameter sweeps without retraining |
| Physical optics model - depth-dependent PSF, crosstalk, staining gradient modeled explicitly | Learns optical artifacts from data; may not generalize to different acquisition conditions |
| Zero GPU requirement - generates on any CPU | Training requires substantial GPU resources; generation is fast after training |
| Encodes only explicitly parameterized biological variation; unknown sources of variation require GAN-based style transfer. | Implicitly captures all biological and technical variation present in training data |
| Parameterization requires domain expertise; LLMs can assist by proposing biologically plausible parameter ranges and guiding initial estimates. | Learns from data without requiring explicit domain knowledge |

## 4 COMPARISON WITH GAN-BASED AND LEARNING-BASED SYNTHESIS

### 4.1 The case for parametric synthesis

The dominant paradigm for photorealistic synthetic image generation is generative adversarial networks and, more recently, latent diffusion models. These approaches have produced good results in natural image synthesis and have been applied to cell and tissue microscopy. However, for the specific problem of generating organoid training datasets, they face fundamental limitations that motivate our parametric approach.

### 4.2 The complementary relationship

Parametric and GAN-based synthesis are complementary rather than competing approaches. Our parametric framework generates a large, diverse, exactly-annotated dataset spanning the full biologically plausible parameter space. This dataset can then be used to train a GAN or diffusion model that learns to add the residual biological texture that the parametric model does not capture.

One practically relevant workflow is CycleGAN or CUT (Contrastive Unpaired Translation)-based style transfer from synthetic to real: the parametric generator produces structurally correct images with exact labels, and the style-transfer network translates these to match the appearance of a target real dataset without requiring paired correspondences. The segmentation model then benefits from exact annotation while training on images that are photometrically similar to real acquisitions. This approach is complementary to our framework’s role as a ground-truth generator.

## 5 VALIDATION

### 5.1 Validation strategy

We validate the framework across three biologically distinct experimental setups using a two-part strategy. First, we apply our feature extraction and analysis pipeline to the open-access per-cell data published by Ong et al. (2025) [5], establishing what real organoids of each type actually show for each feature. Second, we generate five synthetic organoids per condition from the calibrated presets, apply the same pipeline, and compare the synthetic results against the real distributions. This apples-to-apples comparison, same pipeline, same features, same statistical framework, allows us to assess not only whether the generator produces internally consistent differences between conditions, but also how closely the synthetic feature distributions reflect published biological measurements. Statistical comparisons use Mann-Whitney U tests with Benjamini-Hochberg FDR correction at the organoid level, treating each organoid as the independent replicate.

For each condition, synthetic organoids are generated from calibrated presets, segmented using Cellpose 3D [4] (nuclei model for DAPI channel, cyto3 model with two-channel input for cell bodies), quality-filtered by volume and nucleus-to-cell ratio, and then processed through a standalone feature extraction pipeline that computes 18 per-cell morphological, intensity, topology, and shape features. Per-organoid medians are used as the unit of observation for statistical comparison, correctly treating each organoid rather than each cell as the independent replicate.

### 5.2 Analysis 1: PDAC spheroids under osmotic stress

We generate five synthetic PDAC cancer spheroids per condition (isotonic control and hypertonic osmotic stress) using the pdac_isotonic and pdac_hypertonic presets. The two presets differ in organoid diameter (170 μm isotonic, 165 μm hypertonic), cell radius (core 6.8/6.7 μm, periph 7.8/7.7 μm), and heterochromatin fraction (0.30 isotonic vs 0.05 hypertonic, encoding heterogeneous open chromatin vs compacted uniform chromatin). Nuclear texture amplitude is identical between conditions (0.38). All other parameters, optics, staining depth, pressure, and packing, are also identical, isolating the cell size and chromatin state differences as the sole sources of variation. Each organoid contains approximately 1,000 to 1,200 cells (isotonic) or 960 to 1,150 cells (hypertonic) after segmentation. For reference, Ong et al. (2025) [5] reported the following per-cell medians from their real PDAC osmotic stress dataset (1,300 isotonic cells, 762 hypertonic cells, 6 fields each): CV_chromatin showed a medium effect size difference between conditions (d=0.663), with hypertonic cells having lower CV consistent with chromatin compaction. Cell volume, elongation, and roundness showed only negligible effect sizes (d < 0.15). We apply the same statistical framework to our synthetic organoids to assess how well the generator captures this pattern.

Per-organoid median features are compared across 18 morphological, intensity, and topology features using Mann-Whitney U with FDR correction. Of 18 features tested, 3 reach FDR significance (pFDR < 0.05). Nuclear volume is significantly higher in isotonic organoids (median 316.3 vs 296.5 μm^3^, d=+3.39), as is medium axis length (7.17 vs 7.10 μm, d=+2.44). Mean neighbor distance is also significant (15.42 vs 15.32 μm, d=+6.88), reflecting the slightly larger cell sizes in isotonic conditions. Cell volume shows the expected direction but does not reach FDR significance (2075 vs 2021 μm^3^, d=+1.89, pFDR=0.20). Minor axis shows d=+2.14 but does not reach FDR significance after correction (pFDR=0.048, ns). These results confirm that the generator correctly reproduces a modest nuclear volume decrease in hypertonic conditions consistent with osmotic water efflux, with preserved cell shape and spatial architecture. Compared with the real Ong et al. data, the synthetic results reproduce the correct direction for cell size (isotonic larger) and correctly show no topology differences. However, the primary discriminator in the real data, CV_chromatin (d=0.663 in Ong et al.), does not reach significance in our synthetic dataset (d=-0.73, pFDR=0.54). The direction is also slightly reversed. This discrepancy is discussed in Section 6.

Cell shape features (elongation, roundedness, prolate ratio, oblate ratio) are not significantly different between conditions (all pFDR > 0.40), which is the expected biological outcome, osmotic stress causes cell shrinkage but does not dramatically alter cell shape anisotropy. Spatial topology features (neighbor count, local density, crystal distance) are identical between conditions (d=0.00), confirming that the packing architecture is preserved and the two conditions differ only in cell size. Radial distribution is also unchanged (d=-0.49, ns), as expected.

### 5.3 Analysis 2: HMECyst spatial topology -- normal versus cyst-forming

We generate five synthetic HMECyst organoids per condition (normal epithelial control and cyst-forming) using the hmecyst_control and hmecyst_cyst presets. The cyst preset uses lumen_fraction=0.78 to create a hollow shell architecture by enforcing a geometric exclusion zone during cell packing and applying centrifugal repulsion during relaxation, ensuring no cells are placed inside the lumen. The control is a solid spheroid with the same cell type parameters. For reference, Ong et al. (2025) [5] reported the following from their real HMECyst dataset (202 control cells from well A01, 255 cyst cells from well A07): crystal distance was 22.7 μm in control vs 17.3 μm in cyst (d=4.41), Ripley ratio was 1.685 vs 2.721 (d=-1.86), n_nuclei_neighbors was 49.7 vs 63.0 at a 50 μm search radius, and nuclear density was 94,869 vs 120,329 nuclei/mm^3^. All three classifiers (logistic regression, Random Forest, XGBoost) achieved AUC = 1.00 on the real data, confirming that topology features cleanly separate the two conditions in real organoids.

Of 18 features tested, 14 reach FDR significance (pFDR < 0.05). The four non-significant features are CV_chromatin (d=-1.53), oblate_ratio (d=-0.55), cell_elongation (d=-2.05), and cell_roundedness (d=+2.05). The strongest discriminators are spatial topology and size features. The largest effect is on minimum neighbor distance, which is shorter in cyst organoids (19.4 vs 22.2 um, d=+14.24), reflecting the tight lateral packing of cells in the shell monolayer. Cyst organoids also have significantly more neighbors within the search radius (median 18 vs 13, d=-8.94), higher local nuclear density (67,143 vs 48,493 nuclei/mm^3^, d=-8.94), and lower crystal distance (24.6 vs 27.4 um, d=+9.77), all consistent with cells packed into a dense shell rather than distributed through a solid volume. The radial position feature independently validates the shell architecture: cyst cells have a median radial_dist_norm of 0.931 vs 0.870 for control (d=-6.78), confirming that cyst cells cluster near the organoid surface.

Cell and nuclear volumes are significantly larger in control organoids (cell volume 10,847 vs 8,397 um^3^, d=+10.7), reflecting the larger cell sizes in the solid spheroid where peripheral cells expand into available space, contrasted with the laterally compressed cells of the cyst shell. Cell shape is marginally different (elongation d=-2.0, roundedness d=+2.1, both FDR significant) but the effect sizes are small in absolute terms.

Logistic regression and Random Forest classifiers trained on the topology feature set both achieve cross-validated AUC = 1.00 at the organoid level, matching the perfect separation seen in the real Ong et al. data. Fig. 5 shows the radial distribution histograms: control cells distribute broadly across all radial zones, while cyst cells are concentrated in a narrow band at radial_dist_norm 0.8-1.0 with no cells in the lumen interior. Fig. 6 shows the topology feature violin plots. The consistency of the AUC = 1.00 result across both real and synthetic data confirms that the hollow lumen architecture is faithfully reproduced and that the topology feature set is sufficient to distinguish solid from cyst-forming organoids regardless of whether the data is real or synthetic.

### 5.4 Analysis 3: Spatial heterogeneity in a large primary PDAC organoid

We generate a single large PDAC organoid using the pdac_large_clustering preset (diameter 200 µm, necrotic_core=true with necrotic_fraction=0.28, apical_elongation=0.12, surface_flattening=0.18, heterochromatin_fraction=0.30). After Cellpose 3D segmentation and quality filtering, 1,090 cells are recovered. This analysis examines whether the generator produces biologically realistic spatial heterogeneity within a single organoid, specifically, a continuous radial gradient of cell properties from the dense necrotic core to the active peripheral layer, without abrupt discrete boundaries. For reference, Ong et al. (2025) [5] reported the following from their real large primary PDAC organoid (3,071 cells): k-means clustering (k=3) produced a silhouette score of 0.185, with intensity features (oblate_ratio η^2^=0.627, minor_axis η^2^=0.625, prolate_ratio η^2^=0.618, average_intensity η^2^=0.588) dominating cluster separation, and topology features secondary (n_nuclei_neighbors η^2^=0.266). Random Forest test accuracy was 0.950.

Radial profile analysis shows smooth monotonic gradients across all major feature categories. Cell volume, axis lengths, and crystal distance all increase progressively from core to periphery. Local nuclear density and neighbor count decrease from core to periphery, reflecting the transition from compressed core cells to the more loosely packed outer layer. These gradients are consistent with the expected biology of large tumor spheroids, where mechanical compression and nutrient gradients drive radial morphological stratification.

k-means clustering (k=3, selected by silhouette analysis over k=2 to 8) produces a silhouette score of 0.174, consistent with the real organoid value of 0.185 reported by Ong et al. Both values confirm that the three clusters exist on a continuum rather than as discrete populations. Despite this low silhouette score, a Random Forest classifier predicts cluster membership with 92.8% cross-validated accuracy (compared to 95.0% in the real data), demonstrating that the gradient is statistically structured and learnable even without hard boundaries.

The top features discriminating clusters by one-way ANOVA eta-squared are intensity and size features: std_intensity_nuclear (η^2^=0.529), cell_volume_um3 (η^2^=0.525), medium_axis_um (η^2^=0.511), avg_intensity_nuclear (η^2^=0.505), and minor_axis_um (η^2^=0.494). CV_chromatin contributes a strong effect (η^2^=0.454), reflecting the combined action of the staining depth gradient, necrotic core bimodal DAPI model, and the bimodal heterochromatin texture. Topology features contribute secondary effects: n_nuclei_neighbors and local_density_per_mm3 (η^2^=0.097 each), crystal_distance_um (η^2^=0.083). Oblate ratio is the only non-significant feature (η^2^=0.003, p=0.25). This feature importance ranking closely matches the real Ong et al. data, where intensity and shape features also dominated (η^2^=0.56-0.63) and topology was secondary. The alignment is notably better than the pre-chromatin-model results, where topology features dominated the synthetic clustering (η^2^=0.65) while intensity effects were minimal (η^2^=0.04). The bimodal heterochromatin model produces the intensity stratification across zones that the previous single-field model could not.

## 6 KNOWN LIMITATIONS

The framework captures the primary morphological and optical features of organoid fluorescence images but does not attempt to model all aspects of real data.

- Cell-cell contacts are approximated by inflated power-diagram boundaries. Real contacts involve specialized junction complexes, adherens junctions, and desmosomes that create locally thickened membrane zones. These are not modeled.
- The necrotic core uses a zone-based approach. Real necrotic zones have continuous gradients from an apoptotic rim to a fully necrotic center, and may contain calcifications, lipid droplets, and extracellular debris affecting optical properties. The three-phenotype model is a simplification of this spectrum.
- Single organoid generation assumes a homogeneous cell population within each zone. Real organoids contain multiple cell types with distinct morphologies that may coexist spatially.
- Mitotic cells (M-phase) are not modeled. These appear as cells with condensed chromosomes and rounded morphology, and are important targets for drug response quantification.
- The optical model assumes an ideal Gaussian PSF and linear photon detection. Real confocal PSFs are not perfectly Gaussian and may have significant sidelobe structure depending on objective NA and immersion medium.
- Extracellular matrix staining is not modeled. Many organoid protocols include laminin, fibronectin, or Matrigel components that contribute fluorescence signal between cells.
- The cyst mode (lumen_fraction > 0) generates cells as a shell but does not enforce a strict single-cell-layer monolayer. Shell thickness is determined by cell size and lumen_fraction. True apical-basal polarity of cyst cells is approximated by high nucleus_ecc_periph rather than modeled mechanistically.
- Within-nucleus chromatin texture uses a two-component model: a sparse heterochromatin blob field (controlled by heterochromatin_fraction) superimposed on a smooth euchromatin background. This produces realistic chromatin granularity and drives strong CV_chromatin stratification within large organoids (η^2^=0.454 in the large PDAC analysis). However, the between-condition CV_chromatin difference for osmotic stress does not reach statistical significance at n=5 organoids per condition (d=-0.73, pFDR=0.54), despite the correct directional intent. The difficulty is that radial fill geometry and staining depth gradients dominate the within-nucleus intensity distribution at the voxel level, and the blob signal contributes a modest modulation on top. Increasing the number of organoids per condition or the blob contrast parameter may improve sensitivity for this specific comparison.

## 7 SOFTWARE AVAILABILITY

The synthetic organoid generator is implemented in Python 3.11 and includes a PyQt5 graphical interface providing a three-panel workspace (Fig 11). The left panel displays a library of all available preset JSON files. The center panel provides a scrollable parameter editor with slider and spinbox controls for all parameters, organized into collapsible sections for Shape, Cells, Optics, Packing, and Texture. The right panel renders a live preview of the organoid at the current parameter settings, updating interactively as parameters are adjusted. A separate viewer tab provides full slice navigation for generated OME-TIFF files with channel mode selection, label overlay, Z-navigation, XY/XZ/YZ views, and PNG export.

**Fig. 1.**
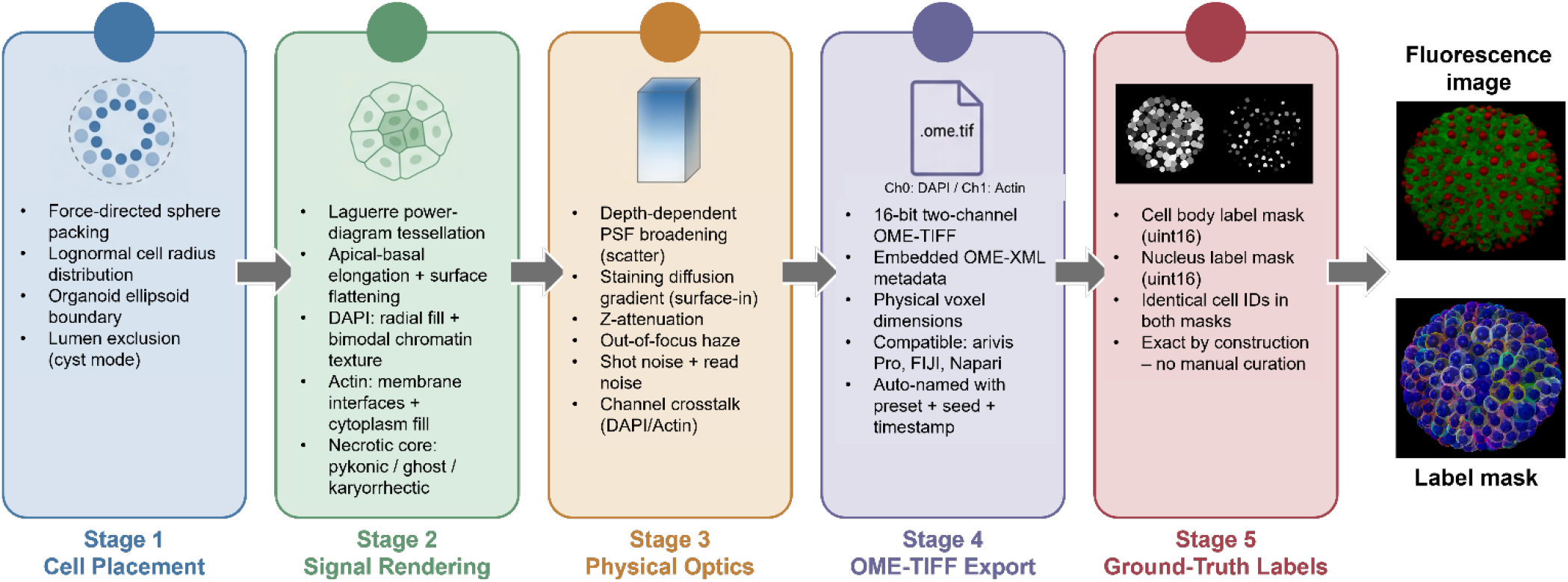
Overview of the synthetic organoid generation pipeline. Five sequential stages produce a two-channel OME-TIFF fluorescence image and accompanying ground-truth label masks.

**Fig. 2.**
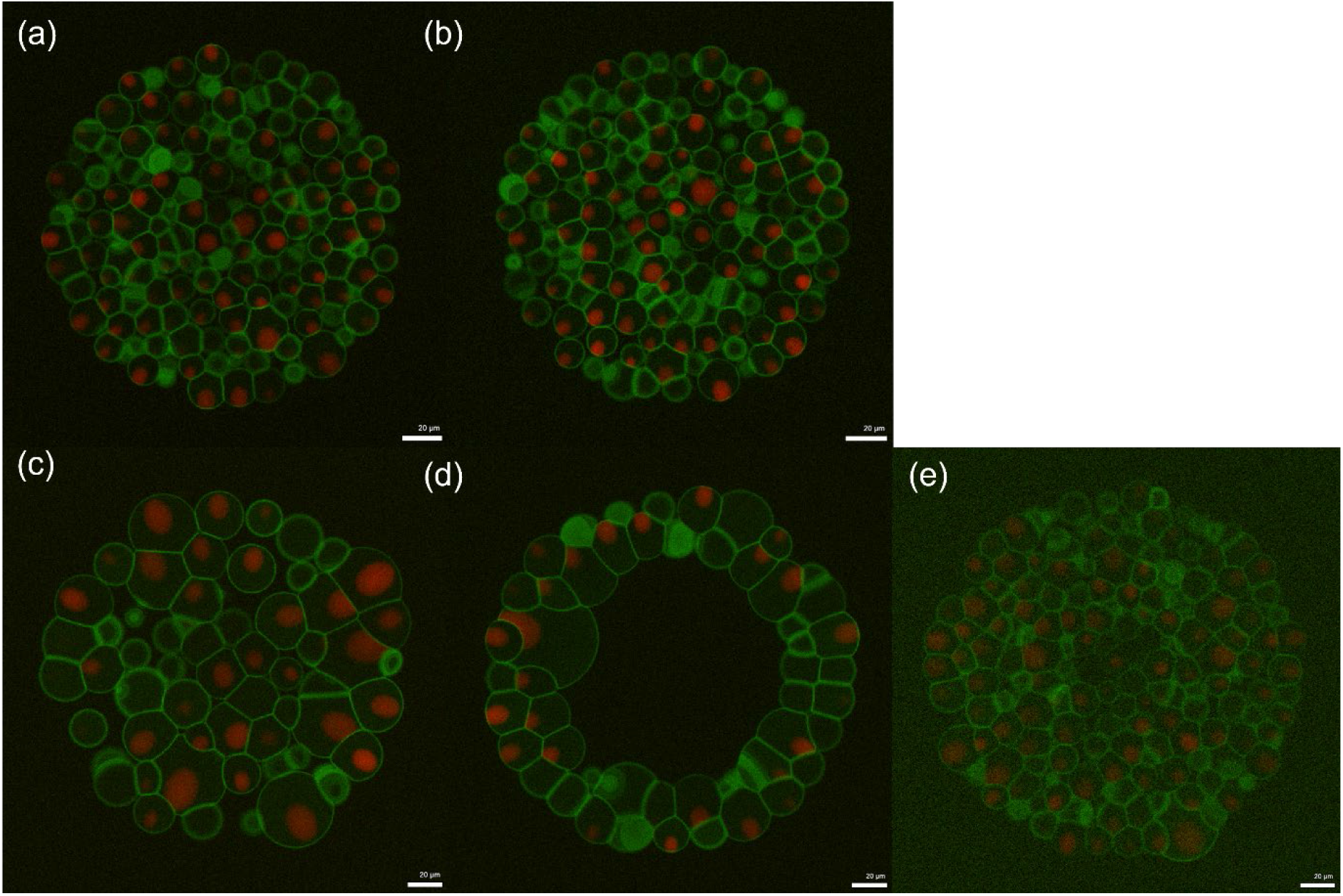
Representative center-plane XY slices of synthetic organoids generated from the five calibrated presets. Nuclear channel in red, actin channel in green. Scale bar: 20 μm. (a) PDAC Isotonic (b) PDAC Hypertonic (c) HME Cyst - Control (d) HME Cyst - Cyst (e) PDAC Large Organoid

**Fig. 3.**
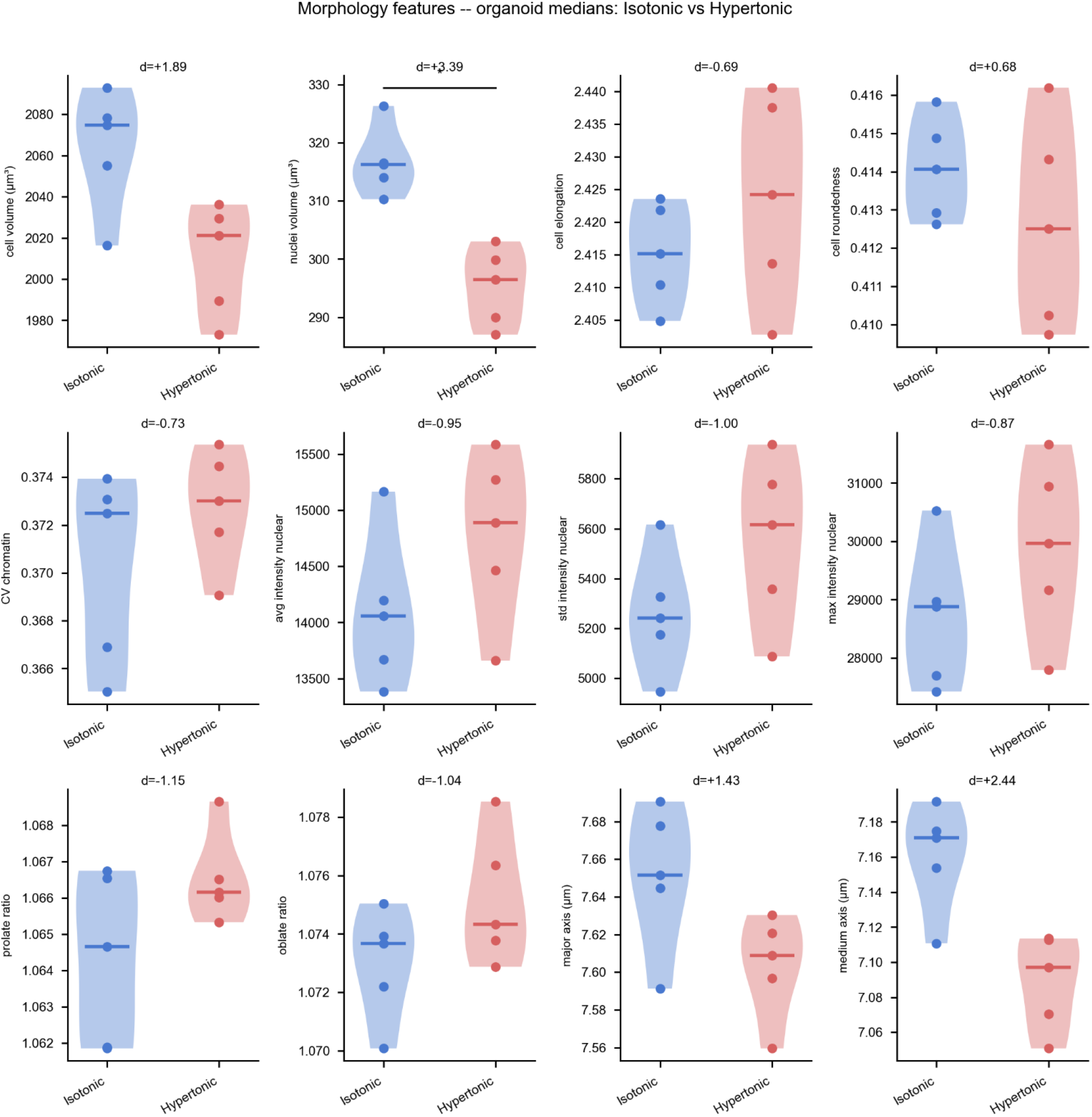
Organoid-level violin plots for 12 morphological features comparing isotonic and hypertonic synthetic PDAC organoids. Features shown: cell volume, nuclei volume, cell elongation, cell roundedness, CV chromatin, avg/std/max intensity nuclear, prolate ratio, oblate ratio, major axis, medium axis.

**Fig. 4.**
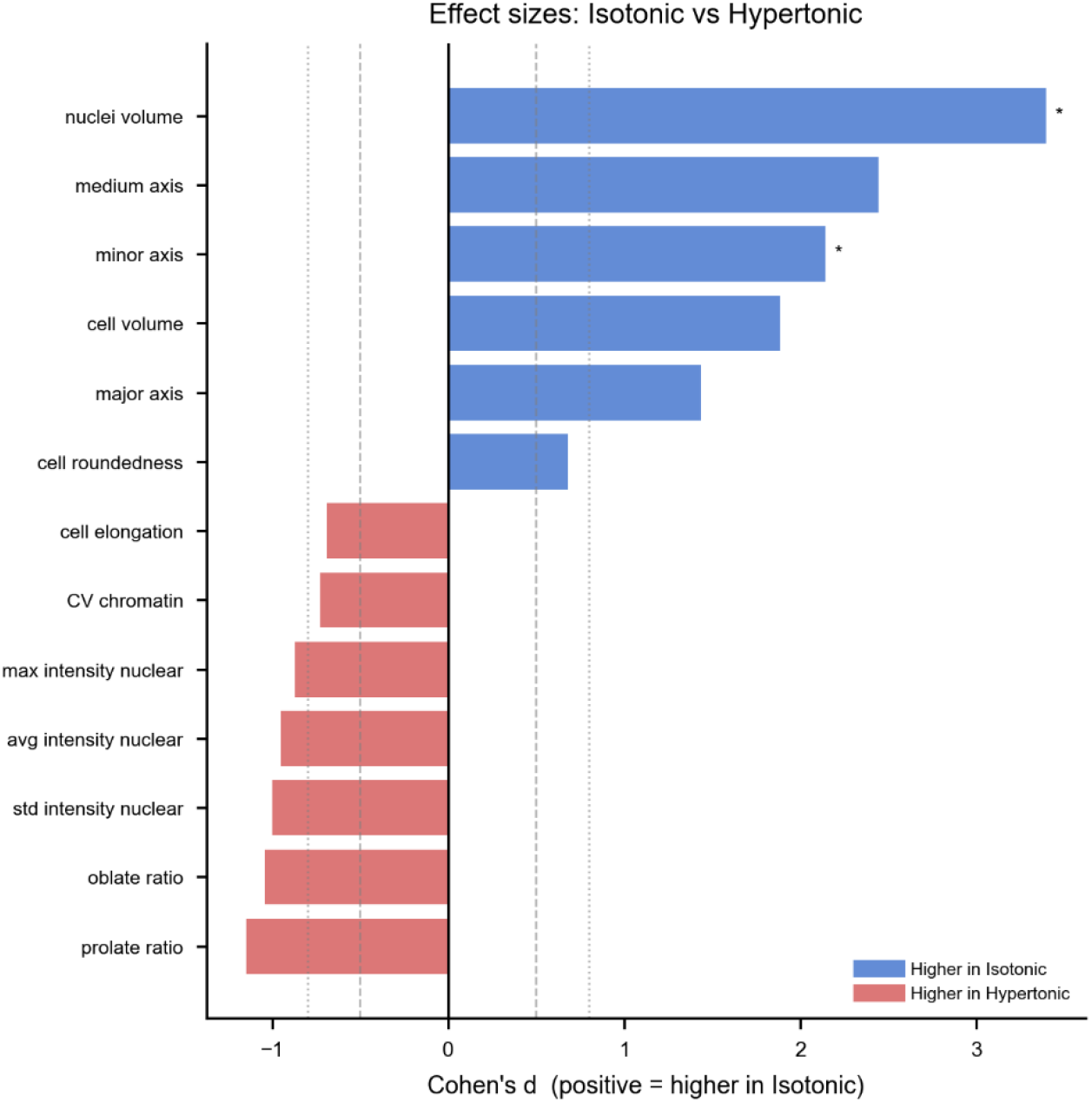
Cohen’s d effect sizes for all 18 features comparing isotonic and hypertonic conditions. Three features reach FDR significance: nuclei volume (d=+3.39), medium axis (d=+2.44), and mean neighbor distance (d=+6.88). Topology features (n_nuclei_neighbors, crystal_distance_um, local_density_per_mm3) show d=0, confirming preserved packing architecture.

**Fig. 5.**
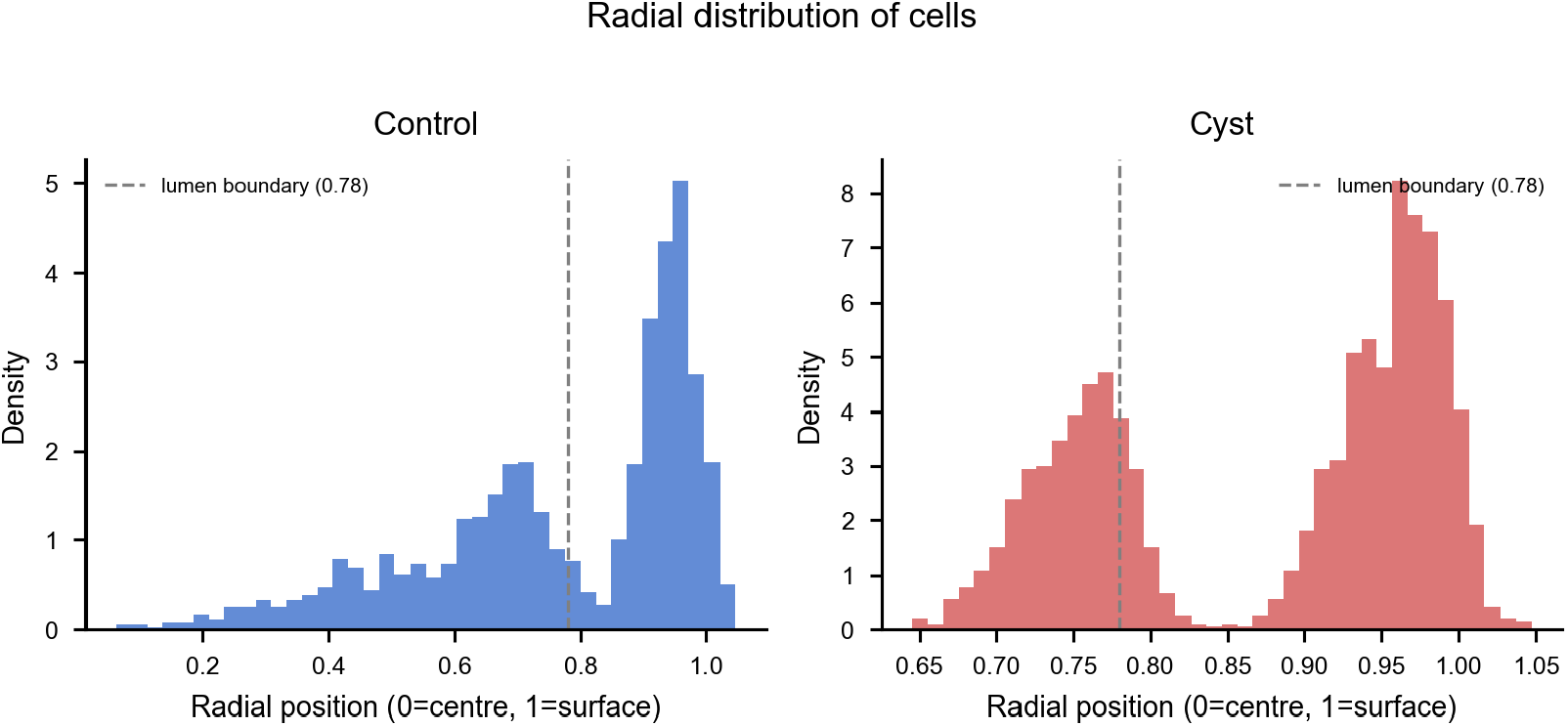
Radial distribution of cell centroids for synthetic HMECyst control (blue) and cyst (red) organoids, pooled across five organoids per condition. Control cells distribute across a wide radial range with density increasing toward the organoid surface. Cyst cells are concentrated in a peripheral shell between radial_dist_norm 0.65 and 1.0, with the distribution peaking near 0.95. The dashed line at 0.78 marks the lumen boundary; in both conditions cells are predominantly peripheral, but the cyst architecture eliminates the broad interior distribution seen in the control.

**Fig. 6.**
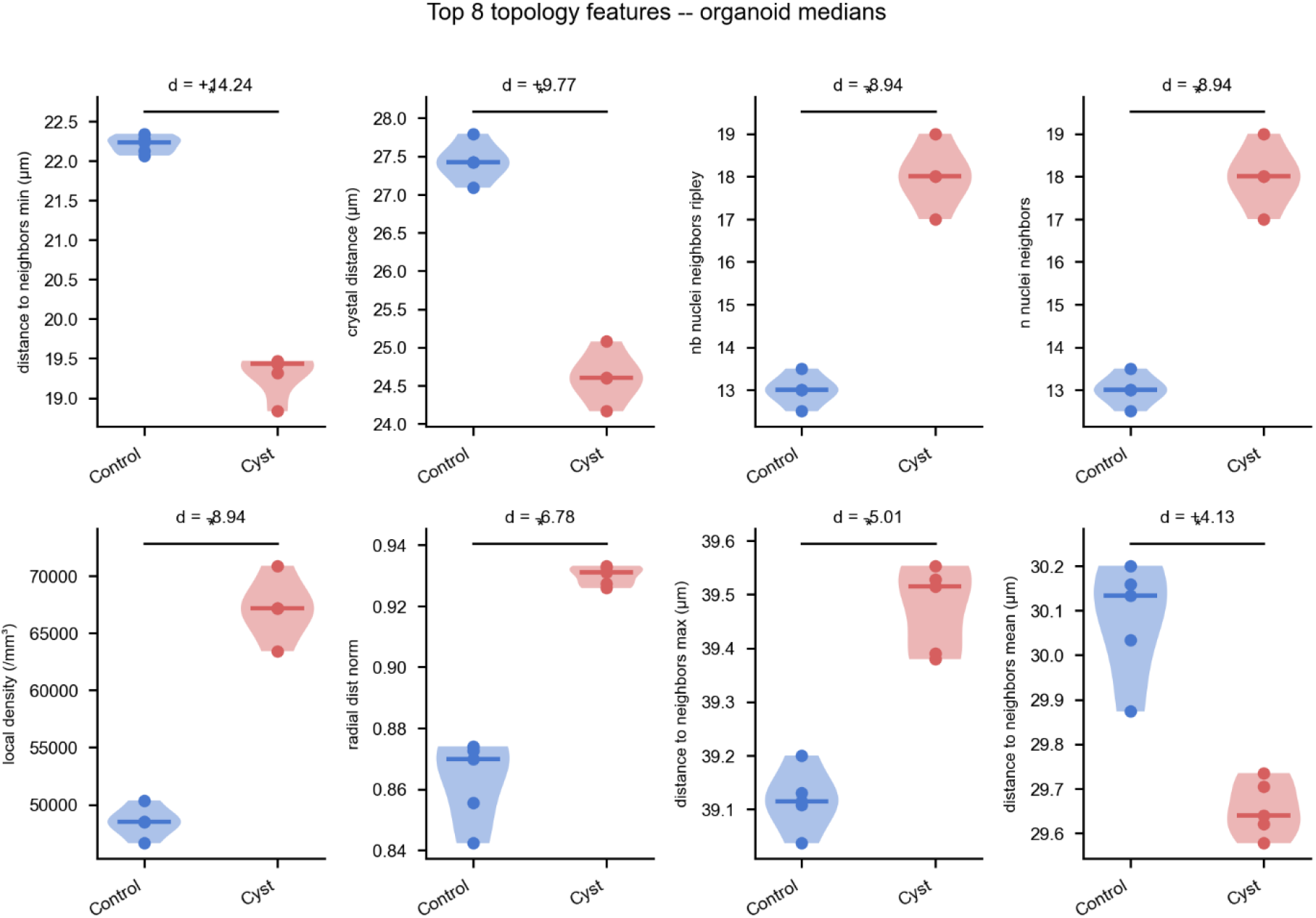
Organoid-level violin plots for the top 8 topology features comparing control and cyst organoids. The strongest discriminators are minimum neighbor distance (d=+14.24), crystal distance (d=+9.77), n_nuclei_neighbors (d=-8.94), local density (d=-8.94), and radial_dist_norm (d=-6.78).

**Fig. 7.**
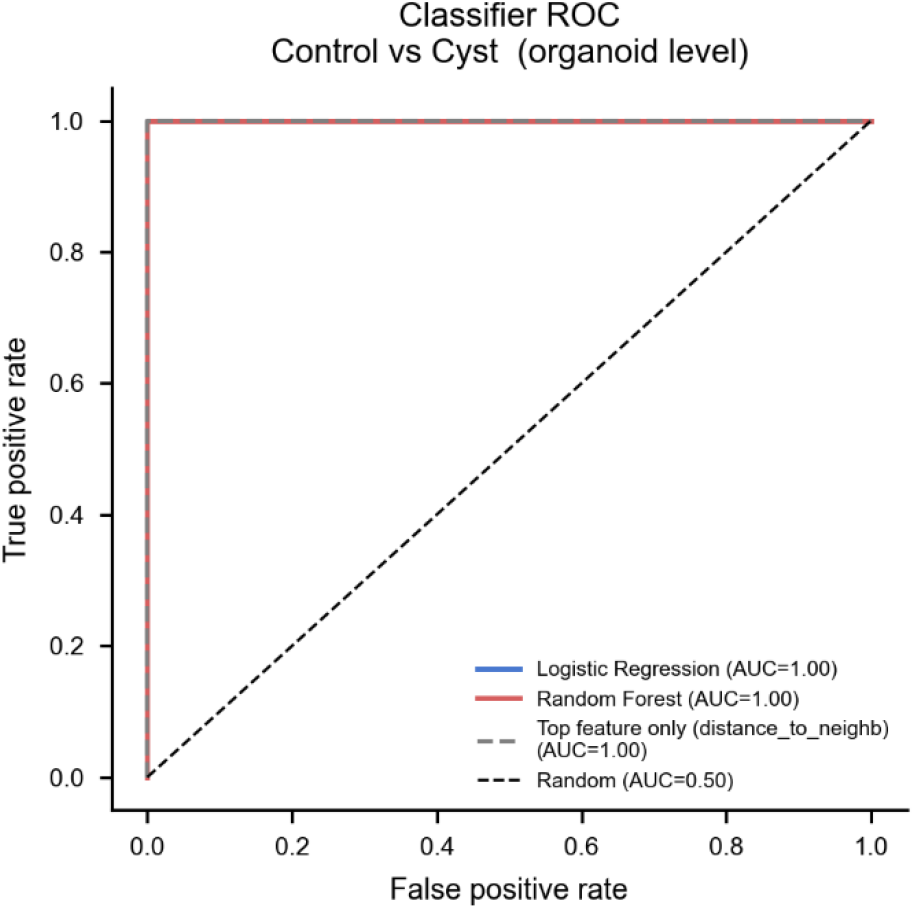
Classifier ROC curves for the HMECyst control vs cyst comparison at the organoid level. Logistic regression and Random Forest both achieve AUC = 1.00. A topology-only feature subset achieves equivalent performance, confirming that spatial architecture alone is sufficient to distinguish the two tissue types.

**Fig. 8.**
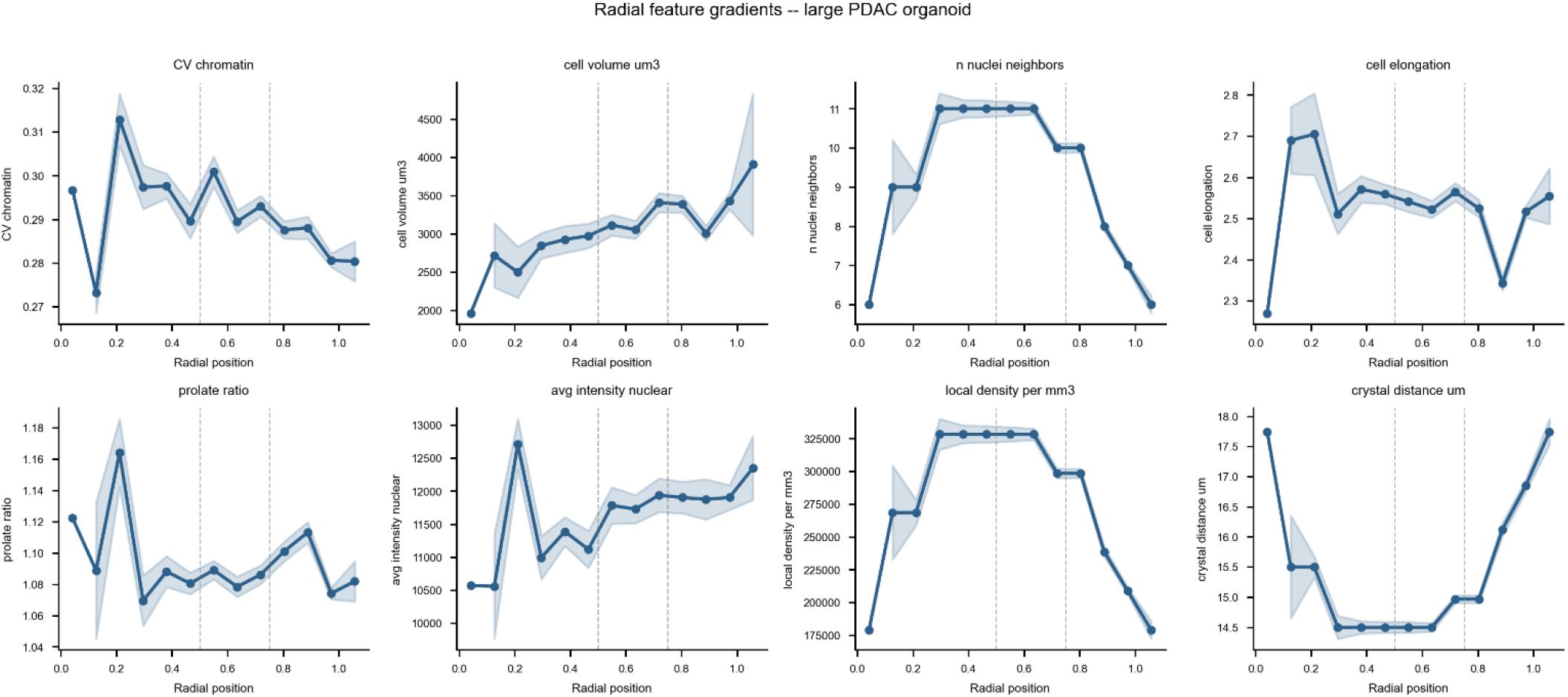
Radial feature profiles for the large synthetic PDAC organoid. Each panel shows median feature value (blue line) with SEM band (shaded) as a function of normalized radial position. Dashed vertical lines at 0.5 and 0.75 demarcate the core/intermediate and intermediate/periphery zone boundaries.

**Fig. 9.**
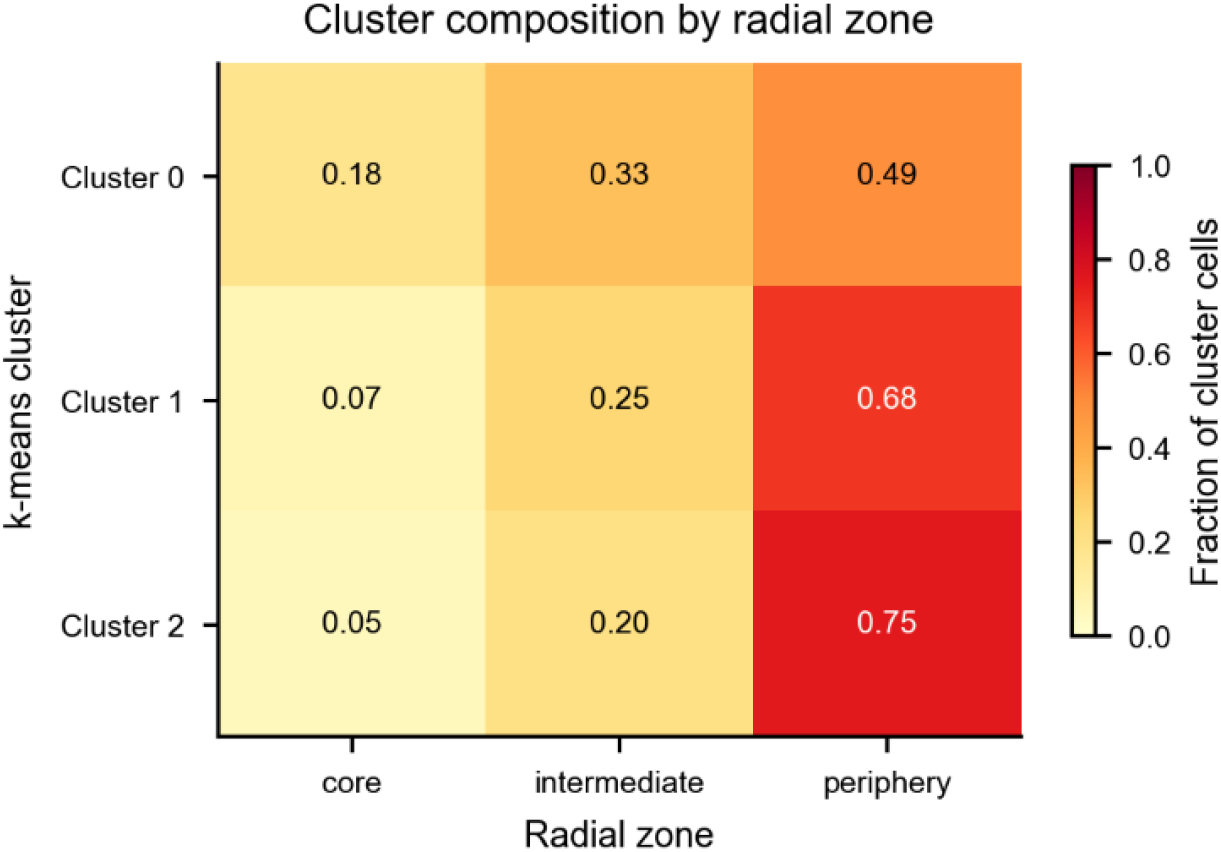
Cluster-zone composition heatmap for the large synthetic PDAC organoid (1,090 cells, k=3). All three clusters are predominantly peripheral: Cluster 2 (75% periphery), Cluster 1 (68% periphery), and Cluster 0 (49% periphery, 18% core), consistent with a continuous radial gradient rather than discrete cell states.

**Fig. 10.**
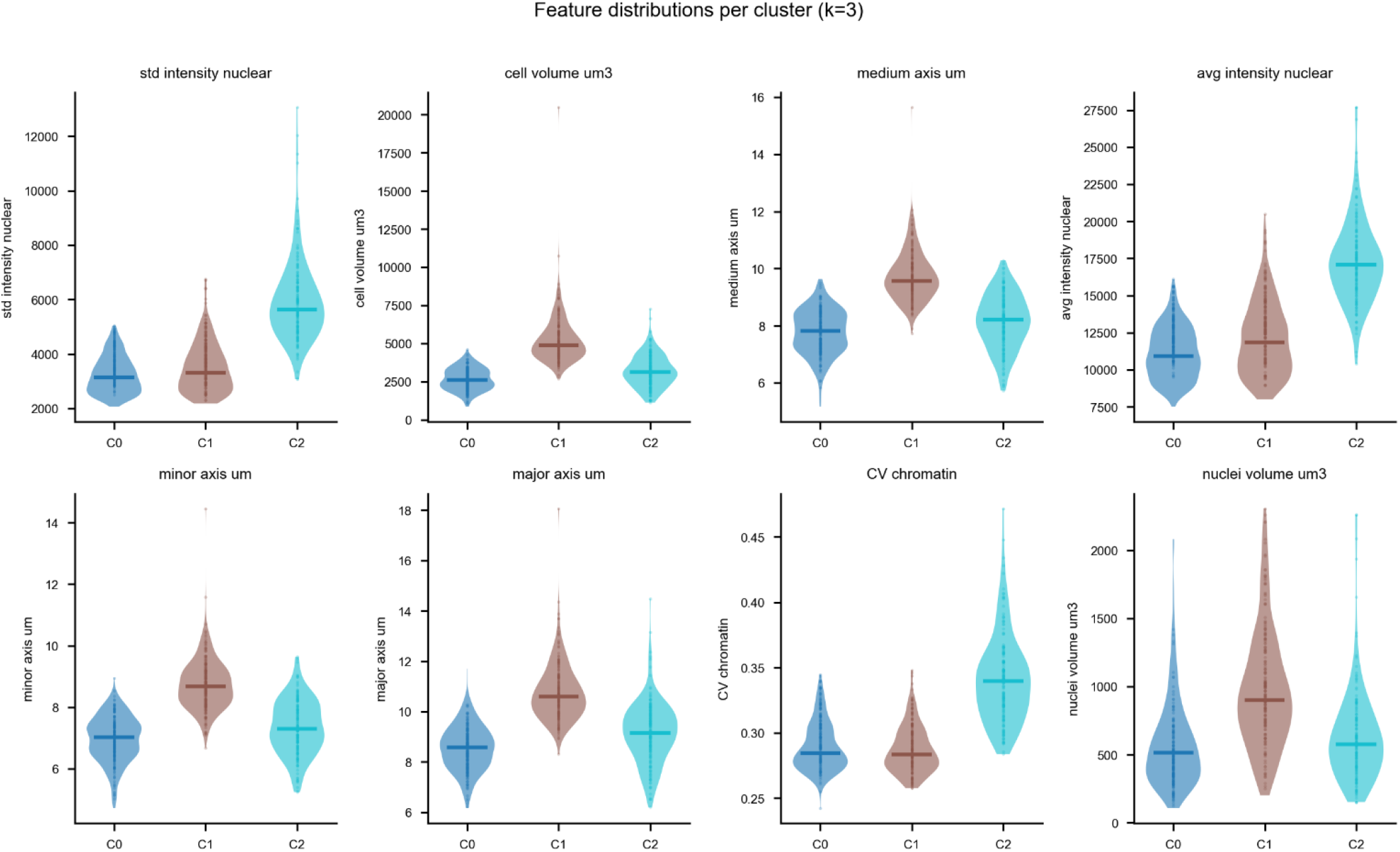
Feature distributions across the three k-means clusters. Intensity features (std_intensity_nuclear, avg_intensity_nuclear, CV_chromatin) and cell size features (cell_volume_um3, medium_axis_um, minor_axis_um) show the strongest cluster separation, consistent with the ANOVA eta-squared rankings. Topology features (n_nuclei_neighbors, local_density_per_mm3) show secondary but significant gradients.

**Fig. 11.**
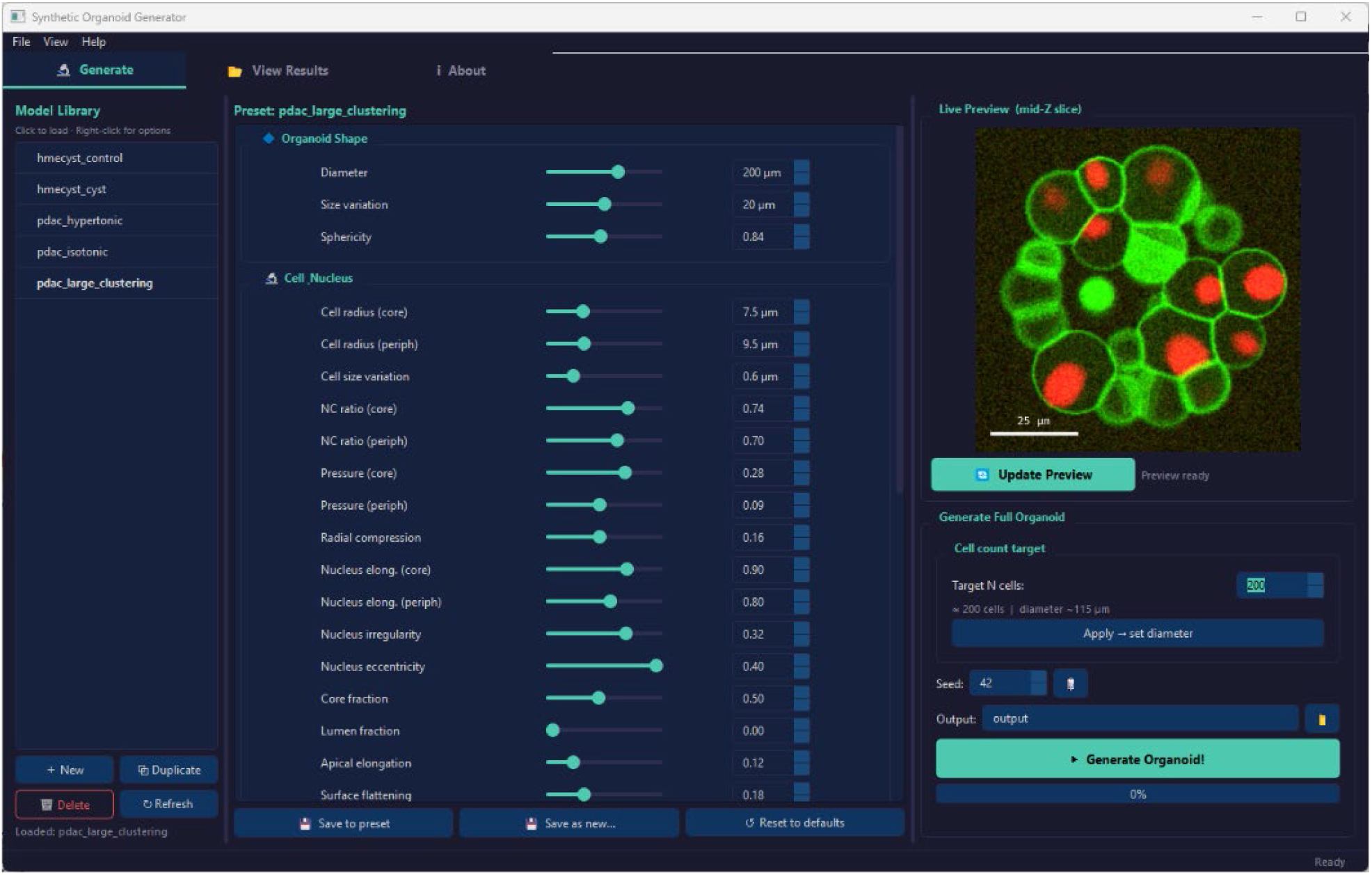
PyQt5 graphical interface for the synthetic organoid generator, showing the preset library (left), parameter editor with sliders (center), and live preview rendering (right).

New organoid types can be defined by creating a JSON file in the presets/ directory specifying any subset of parameters, with unspecified parameters inheriting defaults. A _calibration block is recommended to document the biological measurements and imaging conditions used to derive parameter values.

The source code is released under the GNU General Public License v3.0 (GPLv3), which permits free use, modification, and redistribution for academic and research purposes. Under GPLv3, any derivative work or software incorporating this code must also be released under GPLv3 in its entirety. Organizations seeking to use this software in proprietary or commercial products without these obligations are encouraged to contact the author at to discuss alternative licensing arrangements. The synthetic organoid generator is available at https://github.com/bnsreenu/SynOrg (DOI: 10.5281/zenodo.19616101), and the companion analysis pipeline for segmentation, feature extraction, and topology analysis is available at https://github.com/bnsreenu/SynOrg-Analysis (DOI: 10.5281/zenodo.19616110). Full installation instructions, command-line usage, and worked examples are provided in the README of each repository.

## 8 DISCUSSION

We have presented a parametric physics-based framework for synthetic 3D fluorescence organoid image generation that addresses multiple practical challenges in quantitative organoid imaging: the cost of manual annotation, the absence of ground truth for pipeline benchmarking, the difficulty of stress-testing against rare or complex morphologies, and the need for controlled in silico experimentation before committing to biological experiments. The framework combines biologically grounded cell morphology modeling with an explicit physical optics simulation capturing the depth-dependent optical degradation characteristic of thick tissue imaging.

Three aspects of the current implementation represent advances over prior parametric approaches. First, the hollow lumen mode (lumen_fraction parameter) creates geometrically correct cyst and hollow-shell organoid architectures by enforcing a physical exclusion zone in the packing step and applying centrifugal lumen repulsion during relaxation, ensuring that no cells are physically present in the lumen rather than merely suppressing their fluorescence signal. Second, the unified apical-basal polarity model encodes cell-body elongation and surface flattening directly in the power-diagram distance metric, producing columnar epithelial morphology in peripheral cells and squamous-like flattening at the organoid surface, both effects ramping smoothly with radial position using a t-squared weight function. Third, the three-phenotype necrotic core model replaces a previous uniform pyknotic approximation with per-cell assignments (pyknotic, ghost, karyorrhectic) seeded deterministically by cell identity, reflecting the histological diversity of real necrotic zones.

The staining diffusion model with a 0.15 residual interior plateau and the smooth nuclear eccentricity direction blend (t-squared weight from random in the core to radially outward at the periphery) are additional refinements improving quantitative morphometric accuracy.

The three validation analyzes compare synthetic organoid feature distributions against the real per-cell data published by Ong et al. (2025) [5], using the same analysis pipeline for both. The comparison reveals where the generator succeeds and where specific, bounded gaps remain. For osmotic stress, the generator correctly reproduces nuclear volume decrease in hypertonic conditions (316.3 vs 296.5 um3, d=+3.39) with fully preserved spatial topology (d=0.00 for all packing features) and unchanged cell shape, consistent with osmotic water efflux causing cell shrinkage without architectural reorganization. Three of 18 features reach FDR significance. In the real Ong et al. data, the same three morphological features also show consistent direction, but CV_chromatin was the primary discriminator (d=0.663). Our synthetic CV_chromatin does not reach significance (d=-0.73), a bounded limitation discussed below. For tissue architecture, the hollow lumen mode produces a cyst organoid whose topology features -- minimum neighbor distance (d=+14.24), neighbor count (d=-8.94), local density (d=-8.94), crystal distance (d=+9.77), and radial position (d=-6.78), cleanly separate it from a solid control, with FDR significance across 14 of 18 features and perfect classifier separation (AUC=1.00). This matches the AUC=1.00 achieved on the real Ong et al. data with the same feature set. The directional consistency of all topology features between synthetic and real data confirms that the hollow lumen architecture is faithfully reproduced. For intra-organoid heterogeneity, the large PDAC preset produces a continuous radial gradient with silhouette score 0.174, closely matching the real value of 0.185 reported by Ong et al. Random Forest accuracy (92.8%) is also comparable to the real data (95.0%). Critically, after the bimodal chromatin model was introduced, intensity and size features now dominate the synthetic cluster separation (std_intensity η^2^=0.529, cell_volume η^2^=0.525, CV_chromatin η^2^=0.454), closely matching the real Ong et al. ranking where intensity and shape features dominated (η^2^=0.56-0.63) and topology was secondary. This alignment was not present before the chromatin model fix, when topology features dominated the synthetic analysis while intensity effects were minimal.

The most significant practical advantage over data-driven synthesis remains the provision of exact, arbitrarily detailed ground truth. Because every cell boundary and nucleus position is a direct output of the mathematical model, ground-truth label masks are pixel-perfect and require no manual curation. These exact labels make the synthetic organoids directly usable for benchmarking segmentation algorithms, training deep learning models, and developing quantitative analysis pipelines before any real organoid data is available.

An important clarification about scope: the calibrated presets and validation analyzes in this work are designed to produce organoids whose feature distributions are biologically plausible for the respective cell type and condition. We are not attempting to reconstruct or replicate any specific image from any published dataset. The synthetic organoids are independent generative outputs, and the validation demonstrates that they behave as simulation tools, producing measurable, interpretable differences between conditions that reflect real biology rather than random variation.

CV_chromatin does not reach significance as a between-condition discriminator for osmotic stress (d=-0.73, pFDR=0.54), despite the bimodal heterochromatin model introducing a genuine difference in chromatin domain density between the two presets (heterochromatin_fraction=0.30 isotonic vs 0.05 hypertonic). The difficulty is that radial fill geometry and the staining depth gradient dominate the within-nucleus intensity distribution at the voxel level, and the blob modulation contributes a modest signal on top that the between-organoid variance at n=5 organoids is sufficient to obscure. In the real Ong et al. data, CV_chromatin has a medium effect size (d=0.663) at the per-cell level across 2,062 cells, a sensitivity that five organoids per condition cannot replicate at the organoid-summary level. This is partly a statistical power issue, not solely a generator limitation. CV_chromatin does contribute strongly within the large PDAC heterogeneity analysis (η^2^=0.454), confirming that the chromatin model produces biologically meaningful intensity variation when the within-organoid gradient is sufficiently large. All other expected osmotic stress effects, nuclear shrinkage, axis shortening, preserved topology, are correctly reproduced.

Looking forward, the most productive applications of this framework will be: (1) pipeline benchmarking and regression testing, using organoids with exact ground truth to objectively compare segmentation and feature extraction tools and to detect when a pipeline update introduces errors; (2) stress-testing against edge cases, deliberately generating dense packing, low signal-to-noise conditions, necrotic cores, and atypical morphologies to identify failure modes before encountering them in real data; (3) ablation studies, systematically varying individual parameters to isolate which aspects of organoid biology most affect pipeline performance; (4) imaging system optimization, passing synthetic organoids through varied optical models to evaluate how acquisition parameters and microscope modalities affect downstream analysis; and (5) domain randomization and style transfer, generating training datasets spanning wide parameter ranges and using CycleGAN or CUT-based approaches to transfer real image appearance onto synthetic structure, producing photorealistic training data without manual annotation.

## 10 DATA AVAILABILITY

Code and data availability are described in the Software Availability section. No new experimental data were generated in this study.

## 11 AUTHOR CONTRIBUTIONS

S.B.: Conceptualization, Methodology, Software, Validation, Formal Analysis, Writing - Original Draft, Writing - Review and Editing.

